# Quorum sensing and capsule expression enable subpopulation evasion of phage killing in *Escherichia coli* ST131: Implications for targeted therapy

**DOI:** 10.64898/2026.04.07.717128

**Authors:** Alba Park de la Torriente, Talal Hossain, Sean P. McAteer, Marianne Keith, Gavin K. Paterson, Alison S. Low, David L. Gally

**Affiliations:** Division of Bacteriology, The Roslin Institute, University of Edinburgh, Edinburgh, United Kingdom; Royal (Dick) School of Veterinary Studies, University of Edinburgh, Edinburgh, United Kingdom

## Abstract

*Escherichia coli* ST131 is a globally disseminated multidrug-resistant lineage frequently associated with recalcitrant urinary tract infections (UTIs) and bacteraemia. While bacteriophages offer a promising alternative treatment to antibiotics, their efficacy is often limited in physiologically relevant conditions in comparison to laboratory media. In this study, we have investigated the mechanisms by which the representative ST131 strain, EC958, evades elimination by a model phage, LUC4. We observed that in the urine environment, EC958 can transiently resist phage infection by a density dependent mechanism and by the production of protective polysaccharides. Based on this understanding, we developed a phage treatment strategy that can sterilise an EC958 culture in urine-based medium, even at high bacterial densities. The rational design of the successful phage therapy strategy utilises a tailored phage cocktail containing phage that encode depolymerase enzymes to degrade bacterial surface carbohydrates and the targeting of multiple receptors to prevent the emergence of fixed genetic resistant mutants. We found the addition of specific carbon sources renders the bacteria more susceptible to phage infection. By combining these findings with a simulated bladder wash to model voiding, we successfully achieved elimination of EC958 cultures in a urine environment. This study provides a framework for overcoming both fixed and reversible phage resistance, offering a translatable strategy for effectively treating urinary tract infections with phage.

**Author Summary:** In this study, we investigated how bacterial populations can overcome a phage infection. Phage are viruses that naturally kill bacteria and provide an alternative treatment to antibiotics. We focussed on a particularly aggressive and antibiotic resistant strain of *E. coli*, EC958, which belongs to a group of *E. coli* strains that are a leading cause of urinary tract infections and life-threatening bloodstream infections worldwide. We found that in a simulated bladder environment, these bacteria do not rely on genetic mutations to survive but they employ a range of ‘hide and seek’ strategies. We showed that bacteria can coat themselves in a protective layer to block the phage and use social signalling to enter a dormant state when cell density is high. When they are in this sleep-like state the phage cannot successfully infect. To overcome these bacterial defences, we developed a treatment strategy combining effective phage with specific naturally occurring additives, that trick the bacteria into waking up and becoming vulnerable again to phage infection. By also simulating a clinical bladder wash to reduce bacterial numbers and therefore reduce social-signalling, we were able to eliminate the bacterial population. Our findings suggest that by understanding bacterial strategies we can design more effective and personalised phage therapies to treat bacterial infections.

## Introduction

Urinary tract infections (UTIs) represent a significant global health burden, ranking as the third most common infection worldwide and the second most prevalent in high-income countries (1,2). Annually, UTIs affect approximately 400 million people globally, causing a marked reduction in patient quality of life, particularly in recurrent cases and if left unresolved, can escalate into severe clinical complications (2). Critically, UTI’s serve as a precursor to sepsis, a condition affecting around 50 million people globally each year (3). In developed nations, urosepsis mortality rates remain high, ranging from 20 to 40 percent (4).

The pathogeneis of UTI is well-established, typically involving the translocation of uropathogenic *Escherichia coli* (UPEC) from the intestine to the bladder (5,6). *E. coli* is responsible for greater than 80% of all UTIs (7,8). While these infections are routinely managed with oral antibiotics, the rise of multidrug resistant (MDR) strains both in human and companion animal populations has complicated treatment efforts (9–11). In addition, even with high *in vitro* efficacy, antibiotics can fail to eradicate UPEC *in vivo*, as the ability of UPEC to transition into quiescent, intracellular reservoirs or biofilm-like communities facilitates chronic and recurrent infections (12).

Within the broader category of extraintestinal pathogenic *E. coli* (ExPEC), four sequence types (ST), ST131, ST73, ST69 and ST95, account for the majority of clinical isolates (13–15). Of these, ST131 is of particular concern due to its rapid global dissemination and high levels of antimicrobial resistance. ST131 strains have become the most prevalent ExPEC clonal group (16,17) and are the leading cause of *E. coli* sepsis and associated with increased mortality in bloodstream infections of urinary origin (18).

The rise of MDR strains of uropathogens necessitates the development of alternative therapeutic strategies. Bacteriophage (phage) therapy, the use of viruses that specifically target and lyse bacteria, is re-emerging as an alternative treatment option. Although side-lined in western medicine following the advent of antibiotics the current era of antibiotic resistance has sparked a renewed interest in its application, with a recent systematic review highlighting the potential of phage therapy for treating UTIs (19). However, successful implementation depends on a nuanced understanding of phage-bacterium dynamics within the specific *in vivo* physiological environments.

This interaction is defined by billions of years of evolutionary ‘arms race’ in which bacteria have developed sophisticated defence mechanisms to survive predation (20–22), and phage have evolved reciprocal counter-attack strategies (23,24). In natural ecosystems, these dynamics typically favour the co-existence of both populations; however, the clinical goal of phage therapy is the complete eradication of the pathogen. Achieving this requires overcoming the inherent survival strategies of the bacteria, including the formation of phenotypically heterogeneous subpopulations (25,26). Such phenotypic diversity provides functional resilience during host colonisation and environmental stress, often resulting in a proportion of the population less susceptible to phage attack.

In this study we focus on EC958, a well characterised representative ST131 UPEC strain characterised by its CTX-M-15 production and its resistance to fluoroquinolone, aminoglycoside and trimethoprim-sulfamethoxazole antibiotics (27,28). This broad resistance profile makes it an ideal model for developing alternative treatments. Building on our previous work where we generated a dataset from thousands of phage – bacteria interactions in an artificial urine media (29), we observed that phage activity is often reduced in physiological relevant environments compared to standard laboratory media, such as Lysogeny Broth (LB). Furthermore, while certain phage initially prevent EC958 growth in urine, the bacterial population frequently exhibits rapid recovery even when multiple phage are combined in a cocktail. Here we investigate the different ways by which EC958 evades phage predation, initially by studying a single phage-host interaction, to establish a strategy capable of achieving bacterial eradication in a urine environment.

## Results

### Phage activity against *E. coli* ST131 EC958 in LB, artificial urine and pooled urine

The susceptibility of EC958 was evaluated against a panel of broad-host range phage in LB, artificial urine-based medium (AU) and pooled urine (PU). Interactions were scored using growth assays by calculating the ratio of the area under the curve (AUC) of the phage-infected culture to that of the untreated control over 18 hours. Lower scores represent inhibited bacterial growth, with phage scoring below 60 defined as active phage (29). Phage-susceptibility was found to be medium-dependent, and the majority of tested phage exhibited reduced activity in both AU and PU compared to LB (Fig 1A). Only one phage (LUC4) retained high activity in urine, while three others (Chap1, Nea2 and D4) demonstrated borderline efficacy in AU and/or PU. LUC4 was selected for further study as it displayed the most robust bacterial killing of EC958 in the urinary environment. However, despite strong initial impact, the bacterial population quickly recovered in both AU and PU (Fig 1B). A similar rebound was observed in LB when LUC4 was used as a monotherapy. Distinct differences in bacterial recovery emerged when LUC4 was applied as part of a phage cocktail. In LB, a phage cocktail successfully prevented bacterial regrowth; however, in the urine-based media, the population consistently recovered. This indicates that different evasion or resistance mechanisms are operating under these two distinct physiological conditions. Taken together with the overall activity data (Fig 1A), the results highlight the specific challenge of achieving complete eradication of *E. coli* ST131 EC958 in a urinary environment.

**Fig 1:**
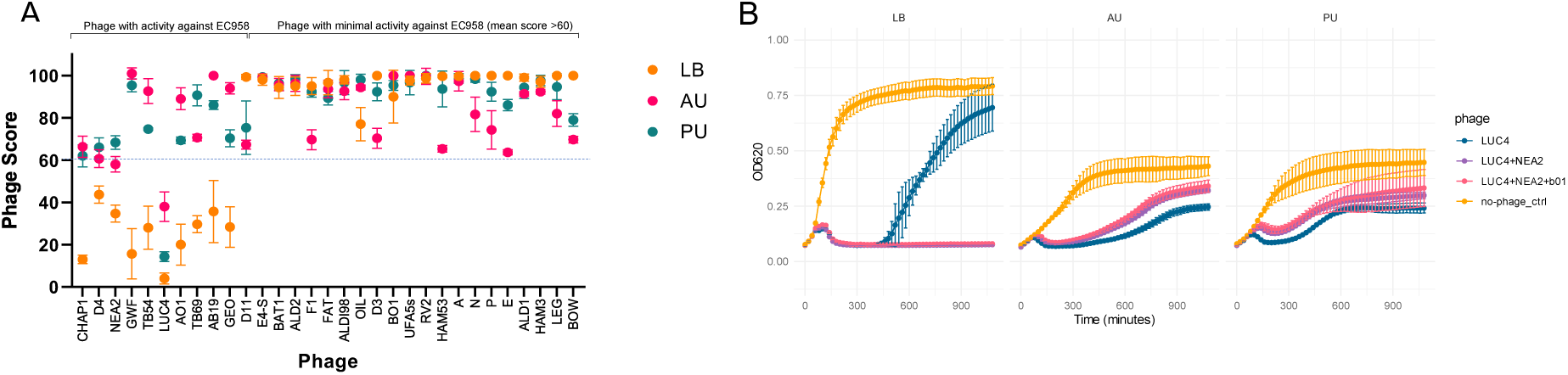
Comparison of phage activity in LB, artificial urine (AU) and pooled human urine (PU). **(A)** Comparison of a set of 31 phage against EC958 in LB, AU and PU. Bacteria were grown in 96-well plates and growth measured in a plate reader. An AUC calculation was used to generate a ‘Phage Score’ with a score of 100 indicating no phage activity and a low score showing high phage activity. A cut off score of 60 was used to define active phage. **(B)** Comparison of single phage and phage combination activity against EC958 in LB, AU and PU over time.

### Characterisation of LUC4

Morphological analysis via electron microscopy confirms that phage LUC4 has the morphology of a myovirus (S1A Fig). Whole genome sequencing further classifies LUC4 within the *Straboviridae* family and the *Tequatrovirus* genus. Its 163,734 bp genome encodes for 49 genes involved in DNA, RNA and nucleotide metabolism, 11 tRNA genes and 10 transcriptional regulation genes. Structural and assembly components are represented by 29 tail genes, 3 connector protein genes and 16 genes involved in head structure genome packaging. Additionally, the genome contains 10 genes associated with host takeover and auxiliary metabolism, alongside 7 genes dedicated to the lysis pathway. 144 genes are still of unknown function (S1B Fig). Bioinformatic screening confirmed the safety profile of LUC4; no virulence factors or AMR genes were detected using the Virulence Factor Database and the Comprehensive Antibiotic Resistance Database respectively. Furthermore, the absence of integration or excision genes indicates that LUC4 is a strictly lytic phage. Physiological characterisation with a one-step growth assay in AU revealed a latent period of approximately 25 minutes and a burst size of ∼78 plaque forming units (PFU) per infected cell (S2 Fig).

### The phage susceptibility of the recovering bacterial populations varies depending on growth media and stage of growth cycle

To investigate the mechanisms allowing EC958 to recover from LUC4 predation in AU, we first examined whether the recovering bacteria exhibited stable genetic resistance or a transient, reversible phenotype. Isolated colonies from the recovered populations were assayed for their susceptibility to LUC4. In LB cultures nearly all recovered colonies exhibited ‘fixed’ resistance, indicative of stable genetic mutations. In contrast, all colonies isolated from the urine cultures after 24 hours were still susceptible to LUC4, indicating a return to the original phenotype (Fig 2A).

**Fig 2:**
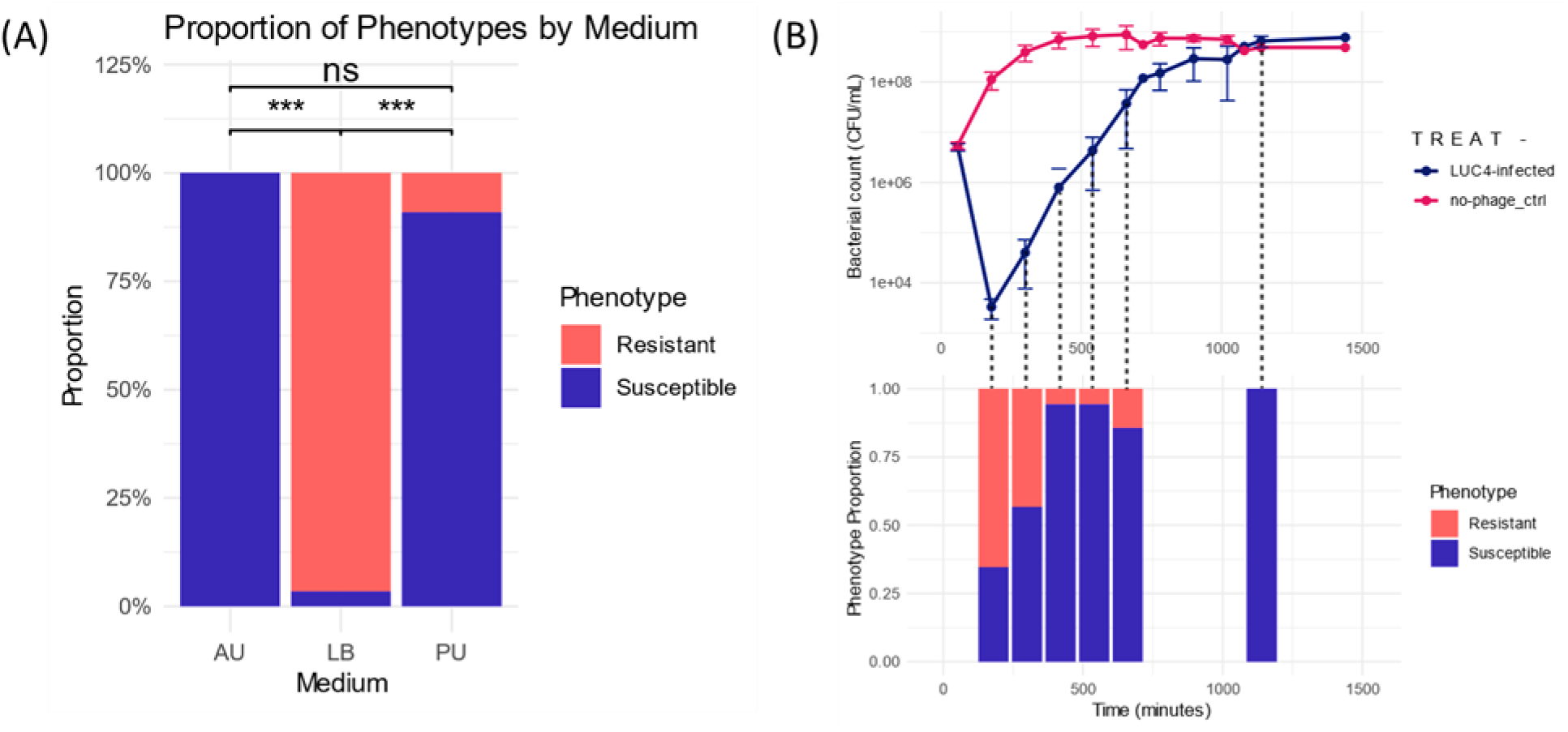
The proportion of recovered EC958 colonies with susceptible and resistant phenotypes to LUC4 by media and timepoint: **(A)** At 24 hours the proportions of recovered colonies with the fixed-resistant phenotype differed significantly between LB and AU (FDR-adjusted p = 3.12e-18), and between LB and PU (FDR-adjusted p = 8.80e-11). In contrast, no significant difference was observed between AU and PU (Chi-squared test with post-hoc pairwise comparisons; FDR-corrected p-values). **(B)** The proportion of recovered susceptible and resistant colonies from AU vary across time in an infection time course: it is mostly dominated by fixed-resistant phenotype in the early timepoints and shifts towards a susceptible dominated population by the end of the assay. The growth curve of EC958 with and without phage represents the mean of 5 biological replicates +_ (error bars).

A detailed time course analysis was performed to monitor phenotypic shifts in population dynamics during the infection cycle in AU (Fig 2B). While a permanently resistant sub-population was detectable and dominant in the early timepoints of the assay, including at the period of maximal phage killing, this was subsequently eclipsed by a transiently resistant subpopulation. This latter group represents the majority of the recovering population, able to grow in the presence of phage in the flask but revert to a susceptible phenotype by the time isolated colonies were assayed. These data suggest that the urinary environment promotes a transient evasion strategy rather than the selection of stable genetic resistance being the predominant phenotype observed in LB.

### Identification of OmpC as the primary LUC4 receptor, its role in genetic fixed resistance and fitness costs caused by its loss

Genomic analysis of 30 colonies from the fixed-resistant populations (from LB and urine cultures) revealed that permanent resistance to LUC4 is primarily driven by the loss of expression or function of the outer membrane porin, OmpC. Genome sequence alignments of permanently resistant bacteria identified a diverse range of structural mutations, including large deletions, insertions and inversions that disrupted either the *ompC* gene itself of its key regulators, *ompR* or *envZ* (S3A Fig, S1 Table).

To definitively identify the LUC4 receptor, *ompC* from EC958 was cloned into the pWKS30 vector and expressed in *E. coli* strain ClearColi®(30), a strain naturally lacking OmpC. While the parent ClearColi® strain was resistant to infection, the induction of *ompC* expression restored full susceptibility to phage LUC4 in both solid and liquid assays (S3B Fig).

The emergence of this fixed resistance, however, is accompanied by a significant biological cost. Competition assays showed that fixed resistant variants (ECRA304.8 and ECRA544.4) exhibited a marked growth disadvantage, with a relative fitness index of <0.5 compared to wildtype EC958 after 24 hours (S3C Fig). These findings suggest that while genetic resistance via OmpC loss provides absolute protection against LUC4, it significantly impairs the competitive fitness of the bacterial population.

Nonetheless, an effective cocktail should include multiple phage that use different receptors to prevent fixed receptor mutants emerging if conditions allow. Therefore, using a LUC4 fixed resistant EC958 OmpC mutant we isolated a new phage, Chap1, that was subsequently determined to use OmpA as its receptor. Chap1 also shows good activity in the urine environment although the bacterial population still rapidly recovers when it is used alone or in combination with LUC4 (S3D Fig and Fig 4B).

### Transient resistance is driven by extracellular factors in the urinary environment

The recovery of ‘susceptible’ colonies following phage challenge in AU is consistent with EC958 expressing reversible evasion mechanism(s) to survive phage predation. This transient resistant subpopulation shows a competitive advantage over fixed resistant mutants lacking OmpC, within the urinary environment. To characterise the dynamics of this evasion, we assayed phage activity at hourly intervals across the growth curve using a constant multiplicity of infection (MOI). These assays revealed that older EC958 populations exhibit increased resistance to phage killing when grown in AU (Fig 3A).

**Fig 3:**
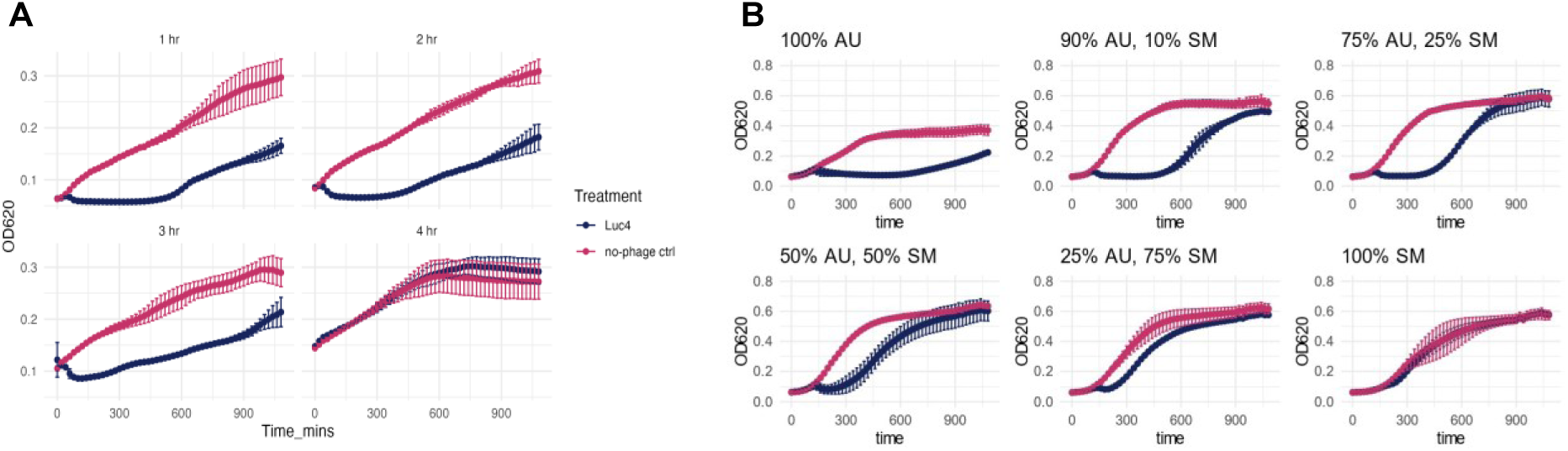
Effect of the media environment on phage activity. **(A)** Aliquots of a growing culture of EC958 were taken at different timepoints (1hr, 2hrs, 3hrs and 4hrs) and challenged with LUC4 (MOI 10) in a 96-well plate growth assay and growth monitored for a further 18 hours. Data shown is from three biological repeats. **(B)** Spent AU medium (SM) from a LUC4-infected EC958 culture (filter sterilised) mixed with fresh artificial urine (AU) in different proportions was used to culture naïve EC958 with phage LUC4. Higher concentrations of spent medium accelerated the development of resistance to phage infection. Data shown is one representative biological replicate, with growth curves displaying the mean ± SD of three technical replicates. Three independent biological replicates were performed.

To determine whether this resistance was a response to the physiological growth phase of the bacteria or to factors accumulating (or depleting) in the environment, we measured phage activity in varying proportions of fresh and spent AU medium (Fig 3B). We found that the spent medium provides protection against phage predation, with bacteria showing maximal sensitivity to LUC4 immediately after dilution into fresh AU. These findings indicate that the transient resistance of older cultures is mediated by external factors. Consequently, we explored two potential adaptive responses associated with environmental sensing that might explain this protective effect: the secretion of exopolysaccharides as a physical barrier and the activation of quorum-sensing-mediated resistance.

### Exopolysaccharides contribute to EC958 evasion of LUC4 phage predation

The secretion of exopolysaccharides, particularly capsule and colonic acid, is a well-established strategy for restricting phage access to surface receptors(31,32). EC958 possesses both a type 2 capsule operon and a colanic acid operon, therefore to investigate their role in evading LUC4 we generated a double deletion mutant by knocking out key genes from these operons (Δ*kpsD,* Δ*wcaF*) (33,34). When challenged with phage one hour post inoculation (approximately 1 x 10^7^ CFU/ml), the mutant exhibited a subtle increase in susceptibility and slightly slower regrowth compared to the wildtype strain (Fig 4A). Further screening across a broader phage panel in AU confirmed this trend, with the double mutant showing generally increased sensitivity to predation (S4A Fig).

**Fig 4:**
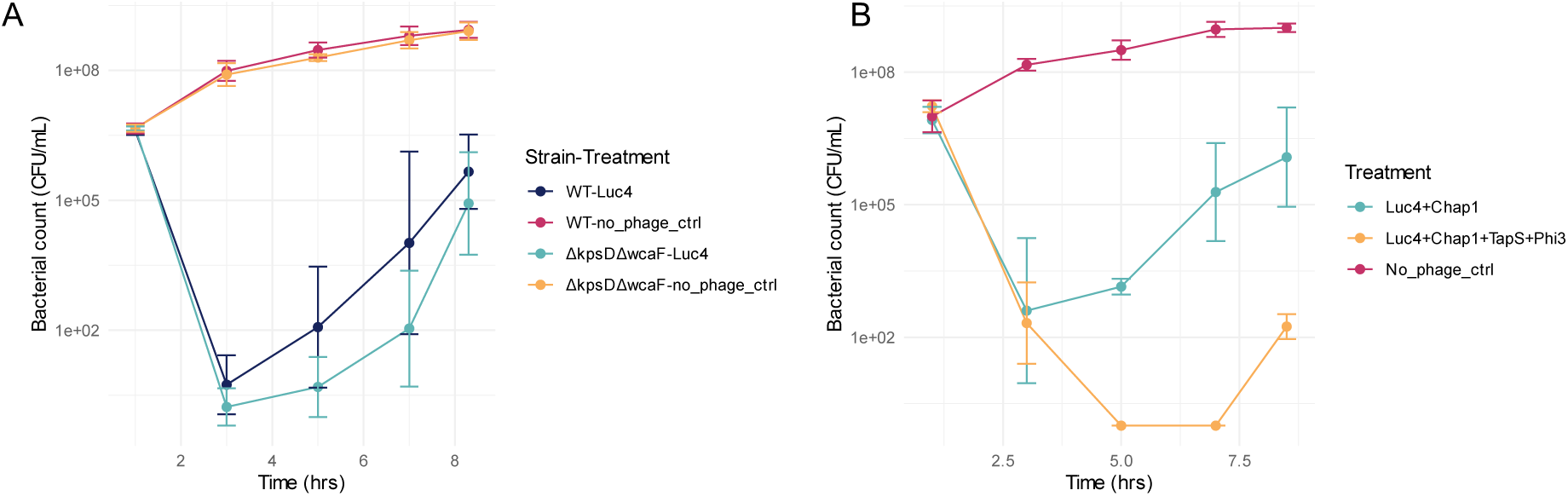
Role of bacterial exopolysaccharide in phage killing efficacy. **(A)** Wildtype EC958 and a double mutant for capsule (CAP) and colonic acid (CA) production (Δ*kpsD*(CAP) Δ*wcaF*(CA)) were challenged with LUC4. Data plotted is from 3 biological repeats **(B)** A comparison of phage cocktails with and without depolymerase active phage. Wildtype EC958 was diluted into fresh AU and either challenged with LUC4 and Chap1 alone or in combination with two depolymerase active phage (TapS and Phi3). Phage were added at MOI=10. Bacterial numbers were enumerated by counting recoverable bacteria on agar plates. Data plotted is from 2 biological repeats.

To overcome this exopolysaccharide-mediated evasion by EC958, we sought to identify phage with depolymerase activity for inclusion in a therapeutic cocktail. We screened for phage against EC958 that produced clear lysis plaques surrounded by expanding turbid halos, an indicator of enzymatic polysaccharide degradation (35,36). This resulted in the isolation of several candidates including TapS and Phi3 (S4B Fig). For comparison, and as a phage with an alternative receptor to LUC4 but lacking visible depolymerase activity we also included Chap1 in this analysis.

Trialling phage combinations against wildtype EC958 showed that a cocktail containing LUC4, Chap1 and the two depolymerase-active phage, TapS and Phi3, increased the efficacy of bacterial killing and further delayed population recovery compared to the combination of LUC4 and Chap1 alone (Fig 4B). However, while the inclusion of exopolysaccharide-degrading phage probably negates the protective effect of surface polysaccharides, a proportion of the population still survives the initial challenge, leading to eventual recovery.

### Quorum sensing and bacterial cell density modulate phage susceptibility

The protective effect observed in high density culture supernatants suggests quorum sensing as a key determinant of phage resistance in EC958. To investigate the role of cell density, an overnight culture was washed and resuspended in AU at densities ranging from 10^1^ to 10^7^ CFU/ml before phage challenge, whilst the MOI of the assays was kept consistent. At starting cell densities at and below 10^5^ CFU/ml we observed effective phage-mediated killing with minimal bacterial recovery (Fig 5A). In contrast, while higher starting densities initially showed phage-mediated killing, the bacterial population consistently recovered.

**Fig 5:**
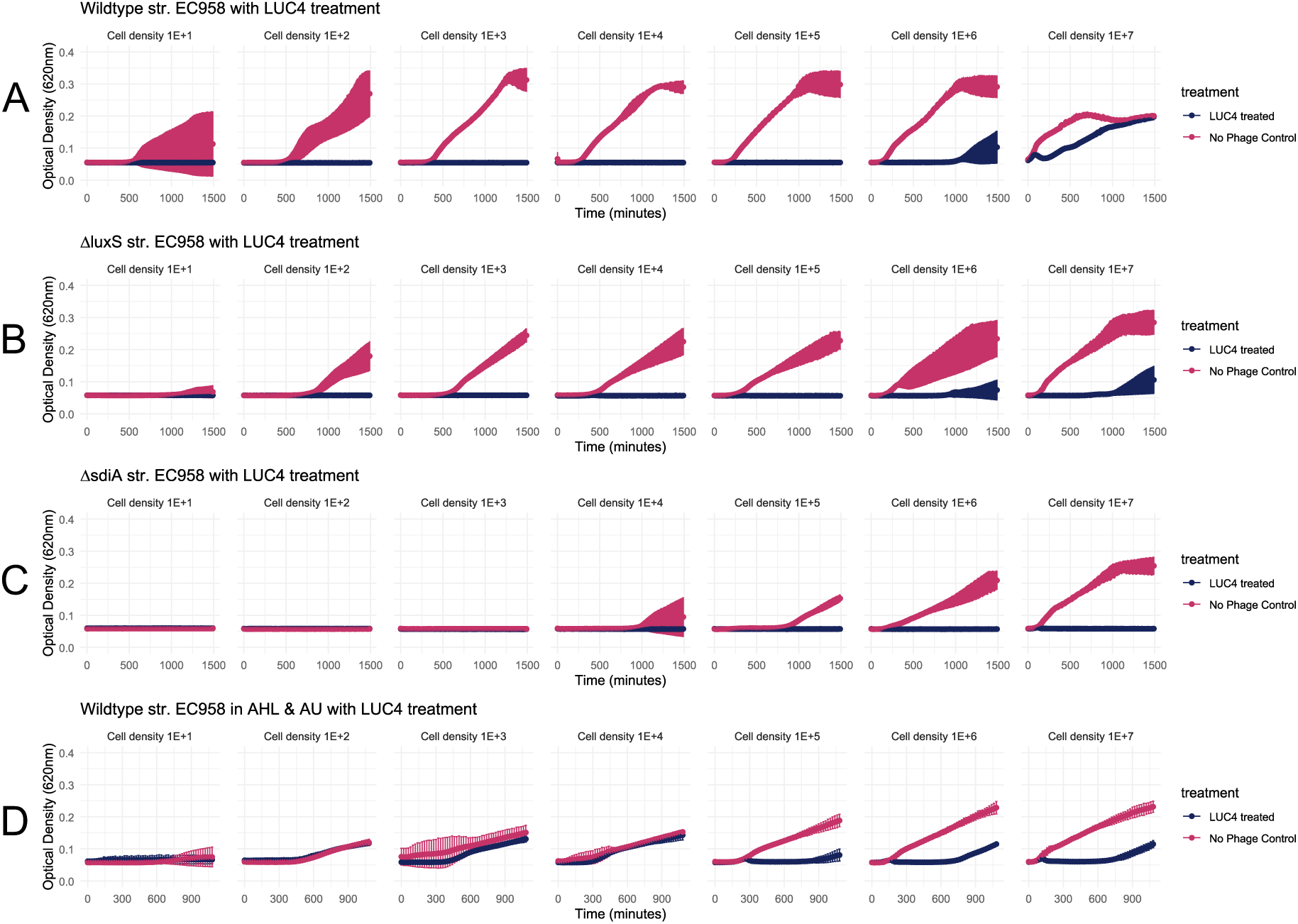
Role of cell density and quorum sensing on LUC4 phage activity against EC958. **(A)** LUC4 phage challenge with different starting densities of wildtype EC958 **(B)** LUC4 phage challenge with different starting densities of *luxS* knockout mutant **(C)** LUC4 phage challenge with different starting densities of *sdiA* knockout mutant **(D)** LUC4 phage challenge with different starting densities of wildtype EC958 in the presence of exogenous AHLs. Data shown is one representative biological replicate, with growth curves displaying the mean ± SD of three technical replicates. Three independent biological replicates were performed.

To identify the specific quorum-sensing systems responsible for the observed recovery phenotype, we deleted two key genes associated with quorum sensing, *luxS* and *sdiA*. Although *luxS* is primarily associated with monitoring interspecies signalling molecules (37), its deletion did increase phage sensitivity (Fig 5B). The deletion of *sdiA* impaired the viability of EC958 at low cell densities, highlighting its importance for density associated survival. At higher starting cell densities, the mutant strain did grow and exhibited increased susceptibility to LUC4 compared to the wildtype strain (Fig 5C). We further validated the role of signalling molecules by demonstrating that exogenous addition of acyl-homoserine lactones (AHLs), autoinducer molecules that are essential in quorum sensing, increased the resistance of the more diluted EC958 populations to LUC4 in AU (Fig 5D). Collectively, these data indicate that quorum sensing plays a major role in the adaptive resistance of EC958. Furthermore, these findings imply that the simple act of bacterial dilution can rapidly transition the population into a more susceptible state.

### Carbon source screening provides evidence of regulatory inputs into phage resistance

We observed that once the EC958 population started to recover from phage challenge, it did so at a rate exceeding what would be expected from a small number of surviving bacterial cells. This observation, coupled with evidence of small colonies on the agar plates after extended incubation, led us to consider the presence of a ‘persister-like’ subpopulation that survives the initial phage killing and then recovers after a pause in growth (38).

To investigate whether the nutritional environment influences this regrowth phenotype, we performed a Phenotypic MicroArray analysis using Biolog PM01 carbon utilisation plate in the presence of phage (39). This exploration revealed many sole carbon sources interfered with LUC4 efficacy against EC958 (S5 Fig) but further investigation of any inhibitory phenotype was beyond the scope of this study. However, a number of carbon sources had a positive impact on LUC4 efficacy including alanine (Ala) and alanine-glycine (Ala-Gly) dipeptide. It is notable that Ala has been described as a resuscitation signal for the recovery of persistent *E. coli* cells (40). To verify the Biolog result, we added Ala-Gly to a set of cell density experiments with EC958 and showed the presence of this dipeptide reduced bacterial recovery at the higher cell densities following LUC4 infection (Fig 6A). While the mechanism of this is not known, it may be stimulating persister-like cells to exit their dormant state, making that mode of resistance less significant in urine-based media. Another notable observation from the Biolog assay was impact of N-acetylglucosamine (NAG) on increasing LUC4 efficacy. When added to the cell density assays NAG, like Ala-Gly, reduced EC958 recovery at the higher cell densities (Fig 6B). This potential for nutrient-driven resuscitation from a paused state likely involves the stringent response and the alarmone (p)ppGpp which is primarily synthesised by RelA (41). A *relA* mutant was generated in EC958, and as with Δ*sdiA*, this proved very difficult to recover at low EC958 starting densities but it did render the bacteria sensitive to LUC4 with no observed population rebound at a starting cell density of up to 10^7^ CFU/ml (Fig 6C).

**Fig 6:**
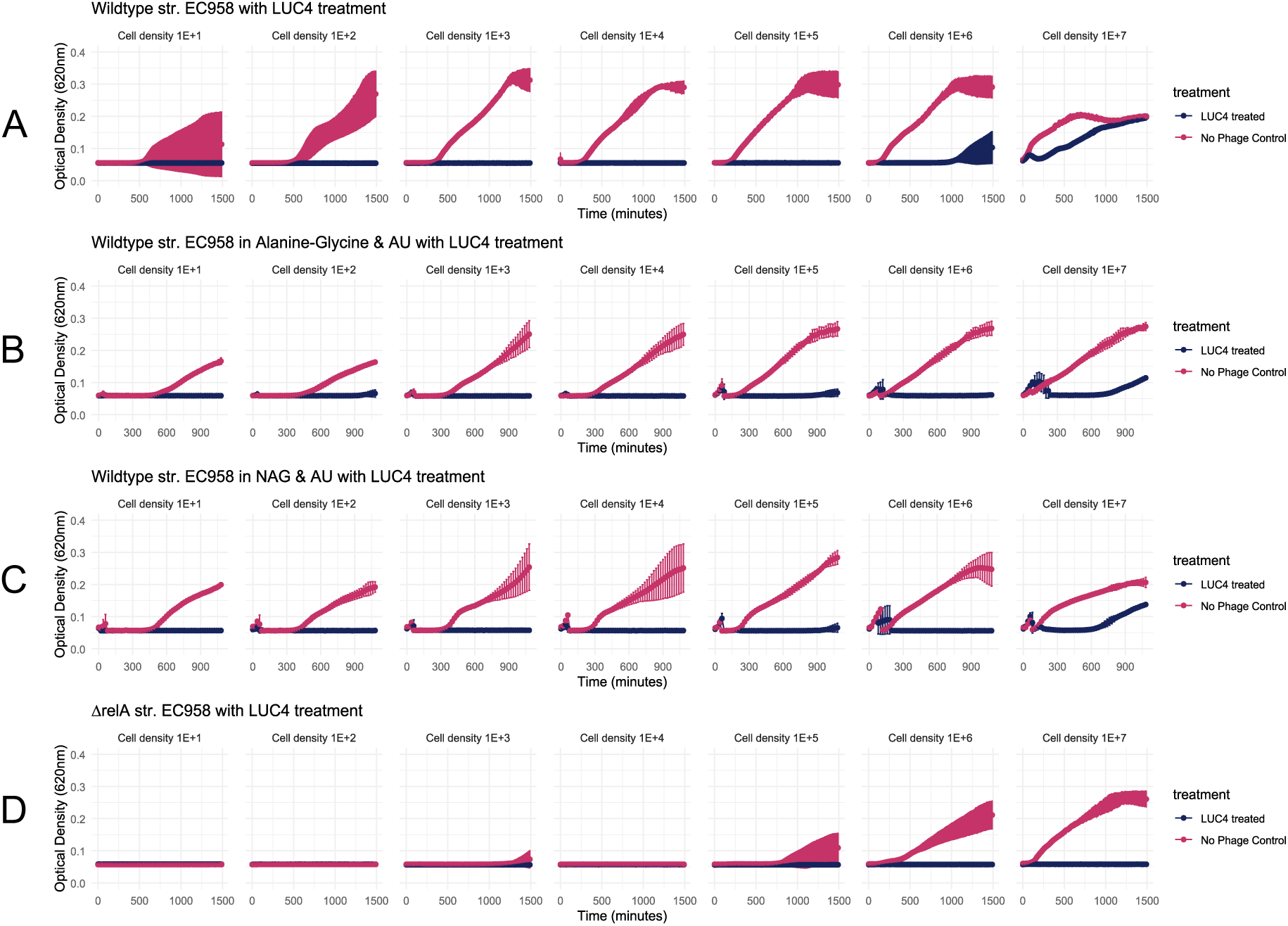
Impact of different carbon sources and the bacterial stringent response on LUC4 efficacy against EC958: **(A)** LUC4 phage challenge with different starting densities of wildtype EC958 **(B)** LUC4 phage challenge of wildtype EC958 in the presence of 20mM Ala-Gly with different starting densities. **(C)** LUC4 phage challenge of wildtype EC958 in the presence of 20mM NAG with different starting densities **(D)** LUC4 challenge of EC958Δ*relA* with different starting cell densities. Data shown is one representative biological replicate, with growth curves displaying the mean ± SD of three technical replicates. Three independent biological replicates were performed.

### Using enhanced understanding of the dynamics of bacterial survival strategies against phage predation to design an effective *in vitro* phage treatment regime

Building on the findings of this study, we have evaluated a four-phage cocktail comprising of LUC4, CHAP1, TapS and Phi3, against EC958 to simultaneously counter exopolysaccharide production and emergence of fixed resistance. This combination targets multiple phage receptors as Chap1 utilises OmpA as its primary receptor while LUC4 utilises OmpC. While the addition of Ala-Gly or potential quorum-sensing inhibitors may enhance bacterial killing at higher cell densities, an alternative straightforward strategy is to minimise bacterial cell density at the point of treatment. In the mammalian host, the bladder regularly voids thereby diluting the bacterial population during a UTI. We modelled a treatment regimen mimicking the catheter-mediated removal of the majority of infected urine (90%), followed by replacement with fresh AU media containing the four-phage cocktail. This strategy effectively eradicated the bacterial population, with no visible recovery of EC958 in the ‘wash’ flasks over 72 hours. To verify this, aliquots were sampled immediately following the addition of the phage cocktail; while the ‘unwashed’ flasks showed bacterial recovery, the ‘wash’ flasks exhibited no growth in 96-well plates (Fig 7). This elimination was further validated by plating the ‘wash’ culture onto LB agar plates at 24 hours; no colonies were observed even after a 48-hour incubation.

**Fig 7:**
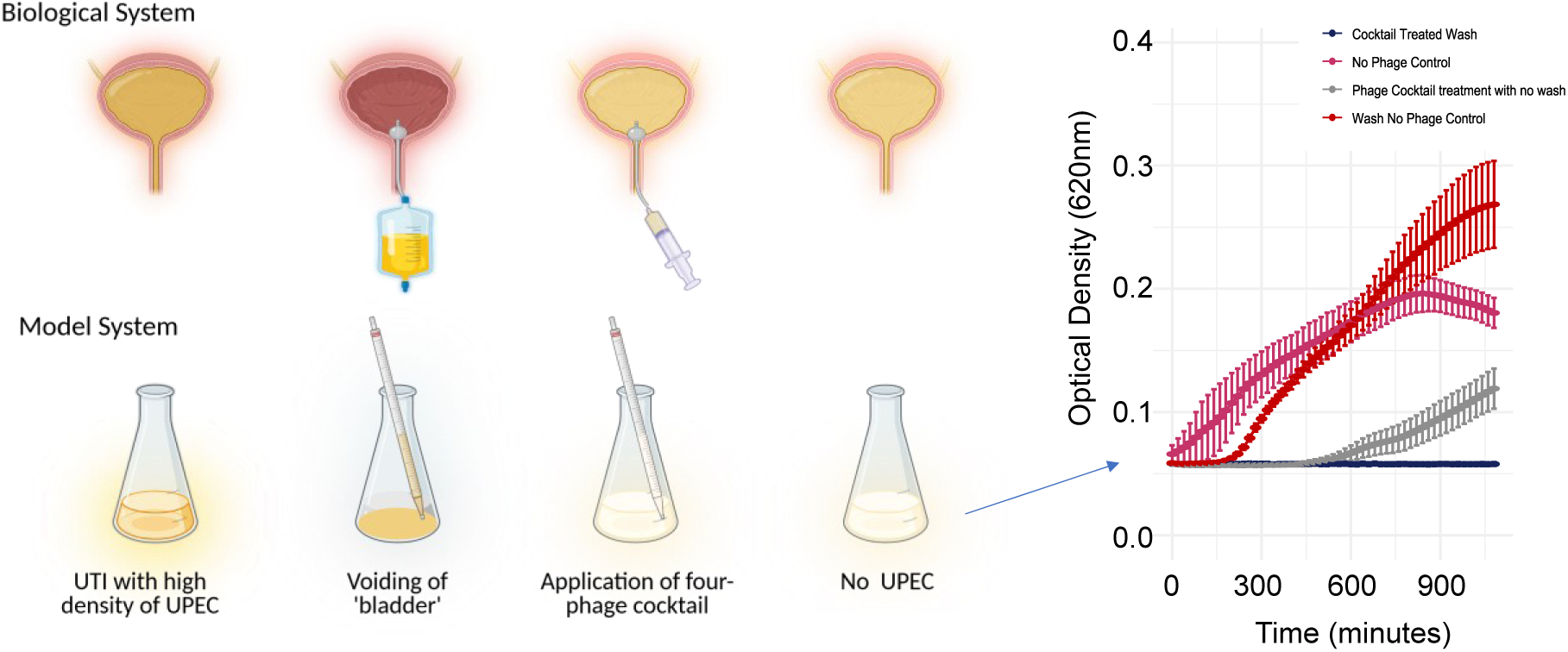
A ‘wash’ mimicking voiding of a bladder just before a phage cocktail treatment prevents EC958 regrowth in culture. 90% of flask contents are removed and the same volume re-introduced with phage cocktail in fresh AU. Aliquots are immediately taken for growth monitoring in multi-well plates and the remaining culture incubated in flasks for 72 hours, visually checking for evidence of growth every 24 hours. 100µl from flask cultures spread onto agar plates at 24 hours and checked for bacterial growth at 48 hours. Experiment was carried out as three independent biological repeats with graph shown as representative data from one biological repeat. Figure created with BioRender.com.

## Discussion

Bacteria and phage co-exist in nature and it is counterproductive for the phage to completely eliminate its host population. For phage therapy to be clinically viable in treating infections such as UTIs, this natural dynamic must be overcome. This study focussed on delineating the interactions between *E. coli* ST131 strain EC958 and LUC4, and the subsequent development of an effective phage cocktail. The initial finding was that LUC4, among other phage, displayed less ability to suppress the bacterial population in artificial urine or pooled urine compared to LB. It was hypothesised that understanding this urine-induced resistance or refraction to infection should facilitate strategies to improve phage therapy for urinary tract infections associated with *E. coli*. The research revealed multiple, potentially linked, levels of resistance from which different strategies were identified to increase phage activity which could then be combined to eradicate *E. coli* in urine-based medium.

A primary level of defence is preventing the phage from adsorbing to its bacterial receptor(s). In a nutrient rich and competition free environment this is often achieved by selecting out a bacterial sub-population that has lost or altered the phage receptor(s). However, *in situ* there are multiple pressures on the bacteria and the mutations will often confer a fitness disadvantage and are therefore less viable long-term strategies during an infection. Nevertheless, the emergence of receptor mediated resistance remains a hurdle. This can be overcome by utilisation of a phage cocktail containing phage that target different receptors. In this study we have demonstrated the many ways that mutations can arise to stop expression of OmpC, the primary receptor for LUC4. By incorporating phage Chap1, which utilises OmpA, the issue of these specific ‘fixed’ mutations being selected was negated.

Our initial studies with LUC4 revealed that the fixed resistant mutants were less competitive in AU medium and were eventually overgrown by a transiently resistant population. The initial hypothesis was that this subpopulation utilises extracellular polysaccharides, such as capsule or colanic acid, to physically shield surface receptors. While capsule expression is a vital UPEC virulence factor, its presents a trade-off; it protects against phage and the host immune system but hinders attachment to host cells (42). Consequently, maintaining heterogeneity in capsule expression is an advantageous survival strategy in the changing host environment. The pathogenicity of UPEC is enhanced by the production of extracellular polysaccharides, such as colonic acid, which facilitates biofilm formation and provides a protective scaffold that shields bacteria from host immune defences and environmental stressors including antibiotics and phage (43).

By utilising a double capsule / colonic acid mutant, we confirmed that these structures confer some degree of protection against phage predation. To counter this in the wildtype strain, we identified phage with depolymerase activity (TapS and Phi3). These enzymes degrade the polysaccharide matrix, removing the bacterial shield. Integrating depolymerase-active phage into the cocktail achieved comparable bacterial killing compared with the non-encapsulated mutant.

Phage with depolymerase activity offer an effective treatment strategy for infections that have formed biofilms and are often more recalcitrant to antibiotic treatment alone (44). Over 80 different capsule types (K-antigen) have been described for *E. coli* and phage specifically targeting some of these capsule types have been described (42,45,46). Since these phage require the capsule as a receptor for infection, their inclusion in a therapeutic cocktail exerts selective pressure that favours reduced capsule expression, ultimately increasing the accessibility and effectiveness of other phage in the cocktail. Identifying specific K-antigens from genomic sequencing and subsequent inclusion of a capsule targeting and/or polysaccharide degrading phage in a phage cocktail should improve treatment efficacy.

However, even with depolymerases a susceptible population eventually recovered, indicating that while surface polysaccharide contributed to transient phage resistance other mechanisms were also present. Our research also implicated bacterial cell density and quorum sensing as critical physiological regulators of phage resistance. Resistance was higher in spent medium and this was validated by mutations in established quorum sensing pathways and addition of AHLs (Fig 5). At high densities, the activation of quorum sensing feedback mechanisms likely serves as an early-warning mechanism to prevent population decimation. Further work is needed to determine the specific quorum sensing mechanisms involved and whether they involve further signalling to phage defence systems in the bacteria.

From the BioLog data it was observed that certain carbon sources, in particular Ala-Gly and NAG could enhance LUC4 activity on EC958. As well as a carbon source, that could signal nutrient rich conditions, NAG has specifically been reported as an inhibitor of quorum sensing (47) and therefore its exogenous addition in our assays could be acting in this way. This offers up different approaches that could be combined with phage to treat the infection; the addition of extra nutrients, the addition of inhibitors of quorum sensing, or rapid urine dilution to reduce cell density at the time of phage addition.

In nutrient rich conditions, such as LB growth medium, EC958 can tolerate receptor mutations and doesn’t appear to display cell density mediated phage resistance. It is clear from the relatively slow growth of *E. coli*, including EC958, in AU that it is a challenging medium and depleted nutrient conditions are a key factor in the formation of a persister subpopulation, which can be triggered by amino acid limitation via the stringent response (48). The presence of persister cells are associated with chronic infections and can impact antibiotic or phage treatment efficacy. RelA is a key enzyme involved in the stringent response to amino acid starvation therefore a *relA* mutant was constructed, and as with the *sdiA* mutant (quorum sensing), it was unable to grow in AU at low population density indicating its critical importance for viability at low cell densities. The response of the quorum sensing and *relA* deletion mutants in our study provide evidence that these systems are contributing to the ability of EC958 to recover after phage infection through transient phage resistance.

While speculative, we postulate that in AU, at higher cell densities (above 10^6^ CFU/ml), a subpopulation of bacteria are present in a persister-like state and would be recalcitrant to phage infection, buying bacteria time to express genes, including phage resistance systems to then allow the bacteria to grow. Whilst counterintuitive, the addition of nutrients to help the bacteria grow could improve the efficacy of a phage treatment. It is proposed that addition of carbon sources such as Ala-Gly or NAG can move the bacteria out of a persister state rendering the bacteria phenotypically sensitive again to the phage. Interestingly, both NAG and Ala-Gly are found in human urine (49) and so their exogenous addition should be easily tolerated. This hypothesis now needs experimental testing but the findings have identified potential adjuvants that could be added to phage treatment formulations to enhance phage activity.

Both the activation of defence systems by quorum sensing and the entering of persister state are closely associated with bacterial cell density. Therefore, if we can manipulate the bacterial cell density when administering a phage treatment then the outcome of the treatment could be greatly improved. The bladder lends itself perfectly for this type of manipulation especially with the use of a catheter for application. The bladder can be emptied, washed and then refilled with phage treatment with the possible inclusion of Ala-Gly and/or NAG to further increase bacterial sensitivity to phage. This, coupled with the use of a well-designed phage cocktail that prevents outgrowth of fixed mutants by including phage targeting multiple receptors and phage with depolymerase activity provides a strategy for effective phage application. In our study, this combination of phage and dilution (single wash and resuspension) was able to prevent any re-emergence of bacteria, even up to 3 days later (Fig 7). Whilst this model system is a closed vessel and missing key *in vivo* components such as cells and immune response the effectiveness of this strategy is promising for future clinical application.

To avoid the fitness costs of genetic mutations, especially in challenging conditions, bacteria employ a variety of non-permanent defences. This research demonstrates that by isolating and addressing each layer of defence individually, we can better understand and ultimately overcome bacterial evasion of phage. The need for an alternative or synergistic treatment for UTI infections is critical with the emergence of MDR UPEC strains that show increased pathogenicity and increased risk of serious complications. This study offers an approach that now needs to be tested across a wider set of key UPEC sequence types and tested in animal or human infections. The efficacy of phage therapy has already been shown to be improved using a personalised approach (50) and the ever decreasing cost and increased speed of whole genome sequencing allows quick identification and genotypic characterisation of an infecting organism including capsule type. In addition, a good understanding of the local epidemiology of circulating clinically relevant bacteria by genomic surveillance has been shown to greatly improve precision phage therapy (51). Analysis of genomic information can now allow rapid treatment decisions as to viability of administering antibiotics or an alternative such as phage. Advances in machine learning models to predict the most appropriate phage to use (29,52), alongside increased understanding of the dynamics of bacteria – phage interactions as demonstrated in this study will pave the way for the clinical use of phage as a treatment for infections.

## Methods

### Strain Information

The *Escherichia coli* O25b:H4-ST131 strain EC958 was kindly supplied by Professor Mathew Upton from Plymouth University. Master stocks were stored at -70°C in 25% glycerol. For use, strains were streaked onto LB agar plates from the frozen stock, and overnight cultures were initiated by inoculating 5 mL of LB, AU, or PU media with a single colony.

*kpsD*, *wcaF*, *luxS*, *sdiA* and *relA* knockout strains were constructed broadly as previously described (53) except using a derivative of pKNG-SceI engineered for chloramphenicol (Cm) selection due to EC958 being kanamycin resistant. This was constructed by amplifying CmR (Table 1) from pBAD33 and cloning into pKNG-SceI using NEBuilder. For Δ*wcfA*, Δ*sdiA* and Δ*relA* flanking regions were amplified from strain EC958 using the primer sets listed in Table 1. For Δ*luxS* and Δ*kpsD* the flanking regions of DNA were generated synthetically by Life Technologies GeneArt (S2 Table). Knockout strains were first verified by PCR using the appropriate -ext primers (Table 1), then confirmed with whole genome sequencing (MicrobesNG, Birmingham).

**Table 1:**
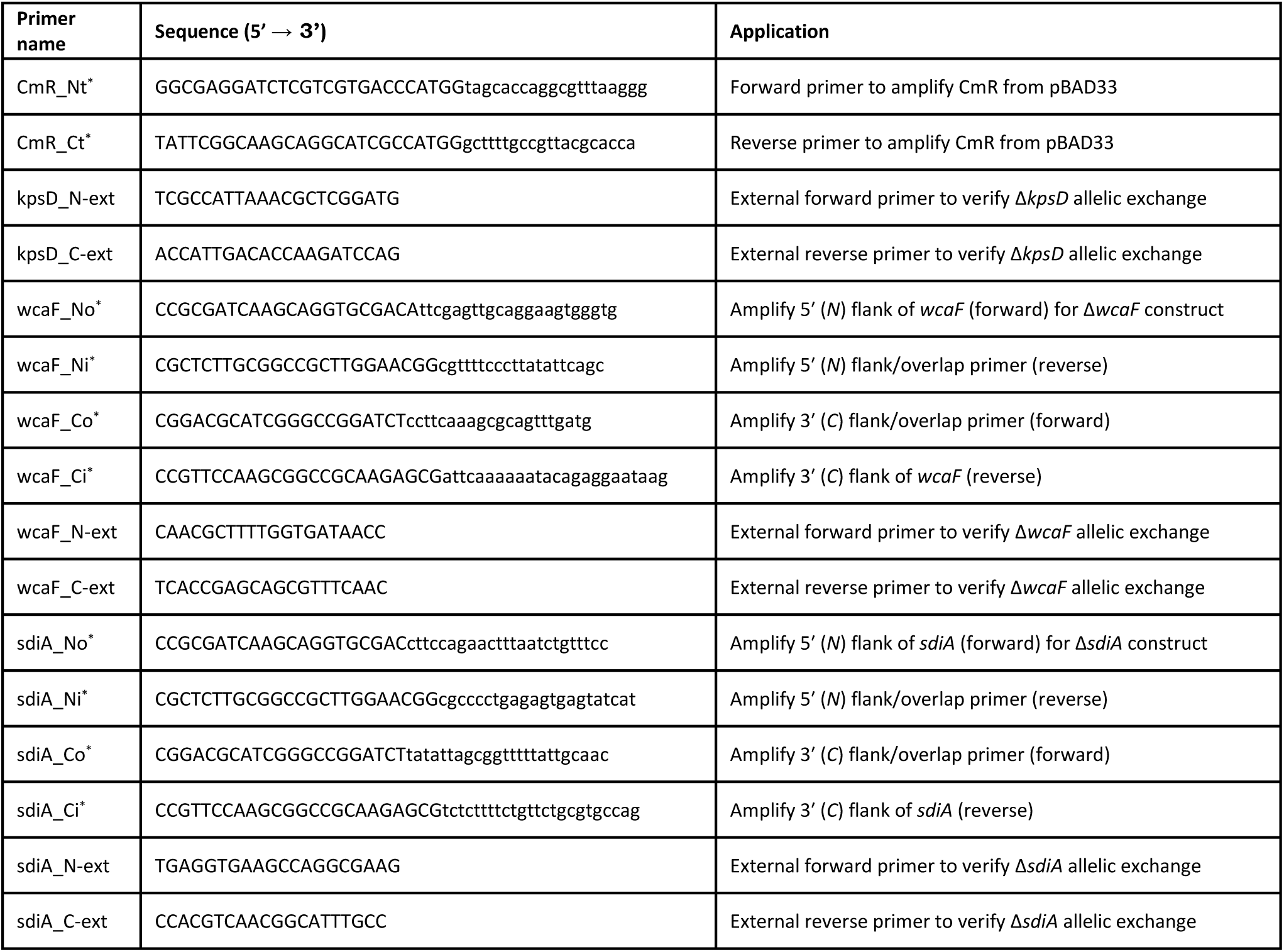
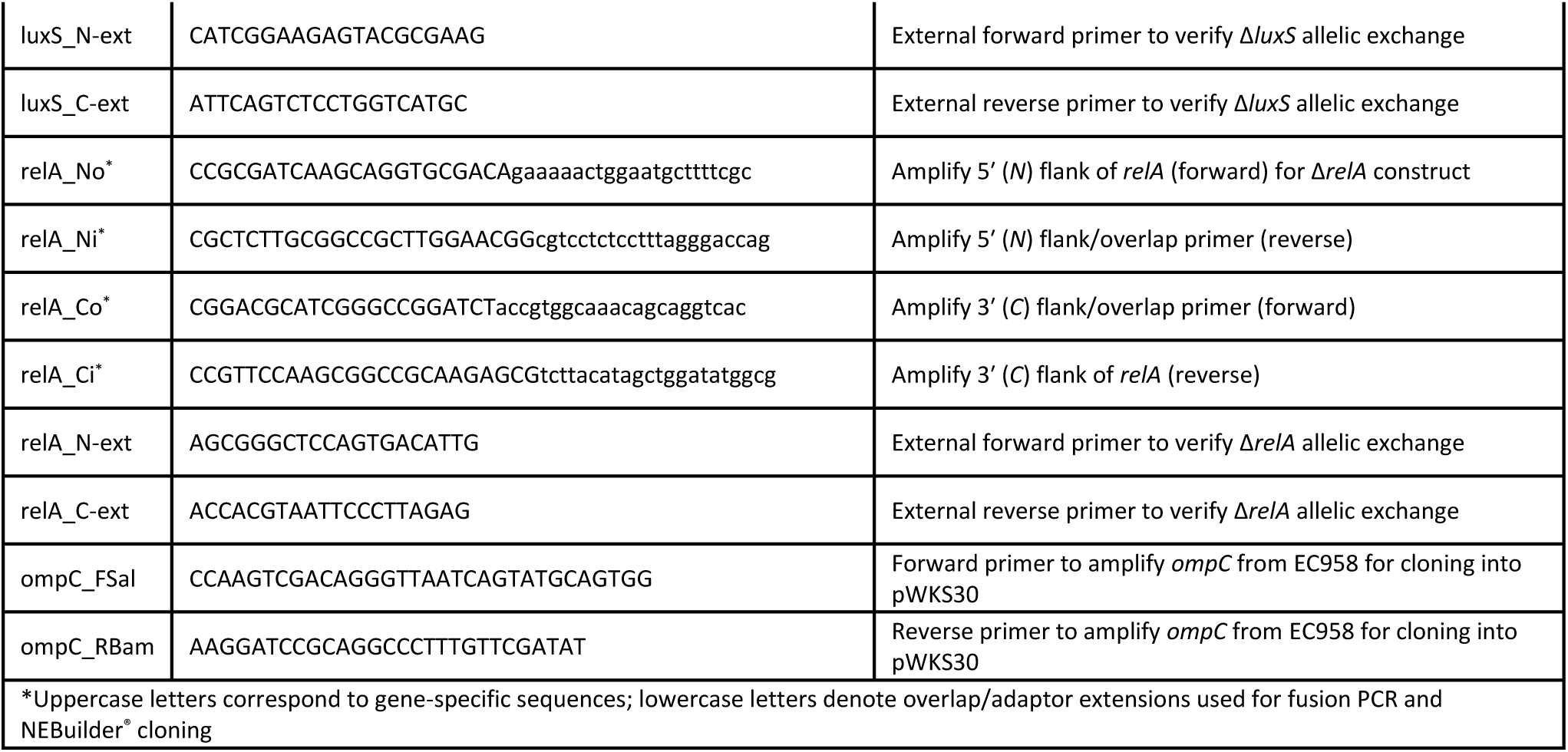
Oligonucleotides used in this study.

### Phage Isolation

LUC4, Chap1, TapS and Phi3 were isolated from wastewater kindly supplied by the Scottish Environment Protection Agency. LUC4, TapS and Phi3 were isolated using wildtype EC958 as a host. Chap1 was isolated using a LUC4 fixed resistant mutant, that was stored after a LUC4 phage challenge. The panel of phage used for Fig 1 and S4A Fig were from a previous study(29).

### Phage propagation

Phages were propagated by subculturing an overnight culture of the propagating host in LB broth at 37°C with shaking for 1 hour; then adding 500 μL of the purified phage and incubating it overnight. 15 mL of the culture was centrifuged at 4,863 x g for 10 minutes to pellet the cells and the supernatant was filtered and retained. Phage stocks were stored at 4°C and diluted into SM buffer [50 mM Tris-HCl pH 7.5, 100 mM NaCl, 8 mM MgSO_4_, and 0.01% gelatin (v/w)] for use.

### Growth Media and Culture Conditions

One litre volumes of AU were prepared (according to the following protocol, https://doi.org/10.17504/protocols.io.kxygx3exzg8j/v1), filter sterilised then aliquoted into single-use tubes, and frozen until required. PU (pooled human urine) was collected from healthy volunteer adults, pooled together into a single batch, filter sterilized using a 0.22-µM syringe filter, aliquoted, and frozen until use.

### Plate reader interaction assays

#### Standard assays

Microtiter plate cultures were prepared by adding ∼10^6^ CFU from an overnight culture (AU and PU) or a starter culture at OD_600_∼1.0 (LB), into microtiter plate wells containing 180 μL of growth media (LB, AU or PU). 10 μL of single phage or phage cocktail (or SM buffer in the no-phage control wells) was added to an MOI of approximately 0.01. Bacterial growth was monitored in a Multiskan FC Photometer (Thermo Fisher Scientific) every 20 minutes over an 18-hour period by measuring absorbance at 620nm, with shaking between measurements. Growth curves were generated with the mean of replicates, and the standard deviation is represented with error bars.

#### Cell density assays

these were carried out as above except the overnight culture was spun down, washed with AU media and the resuspended in fresh AU media and series diluted to give the range of starting cell densities.

#### Calculation of ‘Phage Score’

The mean of three technical replicates was used to create growth curves and calculate the area under the curve (AUC) using PRISM software (GraphPad). [Score = (AUC plus phage/AUC no-phage control) × 100]. For the purposes of plotting the data where the score generated was over 100, it was recorded as a value of 100 representing no phage activity.

### Electron Microscopy

Imaging of phage LUC4 was done with transmission electron microscopy in the Electron Microscope Facilities in King’s Buildings, University of Edinburgh. 20 mL of the neat phage suspension was concentrated using 10,000 MWCO centrifugal concentrator membrane filters (Vivaspin 20, Sartorius Stedim). One volume of the concentrated suspension was diluted in 9 parts of PBS. The preparation of the sample was done by allowing a droplet of suspension to settle on a Carbon 200 mesh Copper grid for 10 minutes and removing the excess solution with filter paper. A drop of 1% aqueous uranyl acetate was applied for 1 minute, the excess was then removed with filter paper. The grids were air dried. Samples were viewed in a JEOL JEM-1400 Plus TEM. Representative images were collected on a GATAN OneView camera.

### Genetic characterisation of phage LUC4

LUC4 genomic DNA was extracted using the Phage DNA Isolation Kit (Norgen Biotek Co. ON, Canada) and Illumina sequenced (MicrobesNG, Birmingham). Reads were assembled with DNASTAR Lasergene and annotated using Pharokka(54) to analyse GC content, antibiotic resistance, and virulence genes. Phage lifestyle was predicted based on the presence/absence of integration and excision annotated genes and using Phage AI(53).

### One-step growth curve

LUC4 host interactions were characterised by a modified one-step growth protocol (54). Briefly, EC958 was sub-cultured in AU and incubated at 37°C for 1 hour. Phage (10^6^ PFU) was added to 9.9 mL of culture and allowed to adsorb for 10 minutes. To synchronise infection, cells were centrifuged (5000 RPM, 10 min, 4°C), and the pellet resuspended in warm AU. To enable direct plating without further processing during the assay, the culture was immediately serially diluted into three pre-warmed AU flasks (A, B, and C). Samples (100 µL) were collected every 5 minutes and plated using the double-layer agar method with EC958. PFU/mL were calculated using the respective dilution factors for each flask. Mean values from three biological replicates were plotted using ggplot2 in RStudio.

### Flask experiments

Overnight cultures were sub-cultured by diluting 1:100 into prewarmed AU in flasks. The cultures were incubated for 60 minutes at 37°C with shaking. Phage were added at an MOI=10 to the infection flask, and SM buffer was added to the control flask. Sampling of the cultures was carried out immediately after the addition of phage and then at regular timepoints. On each timepoint a sample was serially diluted and plated for CFU counts.

### Phage resistance phenotyping

To determine the proportion of phenotypes (susceptible/resistant) of the LUC4 escape population, samples from flask infection cultures were serially diluted and plated on LB agar at specific timepoints. Single colonies were isolated and then purified by passaging them twice on LB agar plates. The phenotype of the isolated bacteria was verified with infection assays in microtiter plate format as described before. The escape variants were classified as ‘resistant’ when they grew similarly to the uninfected control in the presence of phage, and as ‘susceptible’ when they showed phage-driven lysis in the growth curve.

### Genome sequencing

#### Short-read sequencing

Bacterial isolates, including 32 of the variants that showed a fixed-resistant phenotype, were Illumina-sequenced by Microbes NG (Birmingham, UK). The quality of the sequencing was assessed with FastQC.

#### Long-read sequencing of the fixed-resistant variants

DNA was extracted from 10 fixed-resistant variants using the MagAttract HMW DNA Kit (QIAGEN) and purified with the ProNex Size-Selective Puification System, following the manufacturers’ instructions. The DNA concentrations were measured with Qubit dsDNA HS Assay Kit (Invitrogen). DNA samples were barcoded with the Oxford Nanopore Rapid Barcoding Kit (SQK-RBK004), pooled and loaded into a MinION flow cell (Chemistry R9.4.1) following the manufacturer’s protocol. The sequencing was carried out in a GridION Mk1 for a total of 72 hours. The minimum length threshold was set to 200 bp and the minimum quality score to 10. The pores were scanned every 1.5 hours. Base-calling was carried out with the Super-accurate base-calling model of Guppy.

#### Variant calling for the fixed-resistant isolates

The trimmed reads of the short-read sequence were used to search for SNPs or small INDELs using Snippy with the default parameters(57).

#### Alignment and visualization of fixed resistant variants

Porechop (https://github.com/rrwick/Porechop.git) was used to trim the adaptors from the reads. Reads were then filtered by size and quality to a 100x coverage with Filtlong (55) and assembled using Unicycler (59) in the hybrid mode using the Illumina sequence reads described above. Prokka (60) was used for the annotation and Easyfig (61) was used to align and visualise the sequences.

### OmpC receptor characterisation

#### Cloning and expression of *ompC* in ClearColi®

Genomic DNA from EC958 was extracted using the QIAGEN DNeasy Blood & Tissue Kit according to the manufacturer’s protocol. *ompC* was amplified by PCR using Q5 High-Fidelity DNA Polymerase (New England Biolabs) and primers containing *SalI* and *BamHI* restriction sites (Table 1) and cloned into vector pWKS30 to create pEC958ompC. pEC958ompC was electroporated into ClearColi® cells, a strain that does not express OmpC.

#### Infection assays of ClearColi®/pEC958ompC

the cultures were induced by adding IPTG to a final concentration of 1mM, and infection assays with LUC4 were (i) tested in the microtiter plate format as described above, and (ii) in spot assays mixing 100 μL of the induced culture (in log-phase) with 3 mL of molten top agar + 1mM IPTG and spotting a dilution series of LUC4 onto the plates.

### Competition experiments

To demonstrate that fixed-resistant *ompC* mutants are outcompeted by reversible variants, competitive fitness assays were performed. Two *ompC* variants (ECRA304.8 and ECRA544.4) were selected for rifampicin resistance (20 µg/mL) and purified via three rounds of passage. Wild-type EC958 and the rifampicin-resistant variants were co-cultured in 50 mL LB at 37°C for 24 hours. Viable cell counts for each strain were determined at time 0 hours and at 24 hours by selective plating. The fitness index (f.i.) was calculated using the formula: f.i. = LN (Ni (1)/ Ni (0)) / LN (Nj (1)/ Nj (0)) where Ni (0) and Ni(1) represent the initial and final counts of EC958 and Nj (0) and Nj(1) represent the initial and final counts of the resistant variant, respectively. To ensure the observed fitness cost was not an artifact of rifampicin resistance, control competitions were conducted between the parental variants and their rifampicin-resistant derivatives.

### Phenotypic MicroArray (Biolog) Profiling

Metabolic activity was assessed using Biolog PM01 plates (Biolog, Inc., USA) to determine effect of carbon source on phage LUC4 activity. EC958 was cultured overnight on LB agar, and colonies were resuspended in IF-0a inoculating fluid (Biolog) to a target turbidity of 85% transmittance (Biolog turbidimeter). The cell suspension was supplemented with Biolog Redox Dye Mix A. For the phage-challenged group, LUC4 was added at a MOI=0.01. 100 µL was inoculated into each well of the PM01 plate, which contains 95 distinct carbon sources and one negative control (water). Plates were incubated in the OmniLog PM System at 37°C for 48 hours with metabolic activity (tetrazolium dye reduction) recorded every 15 minutes. Kinetic data were analysed using OmniLog PM software. Metabolic curves were compared to identify carbon sources where phage infection was enhanced or inhibited.

### Phage treatment in a model of bladder voiding

This assay followed the previously described flask protocol, with a reduced initial inoculum (1:1000 dilution of overnight culture). Following one hour of growth, a “wash” treatment was performed by replacing 90% of the culture volume with fresh medium containing phage (MOI=10). In “non-washed” treatments, phage was added directly to the existing culture; no-phage controls were processed identically. Straight after phage addition 200µl aliquots from all flasks (in triplicate) were transferred to 96-well plates for 18-hour kinetic analysis. After 24 hours, flasks were visually inspected for clarity, and 100µl aliquots were plated onto agar plates and incubated for 72 hours, with daily monitoring for any delayed bacterial recovery.

## Acknowledgements

Electron microscopy of LUC4 was carried out with the help of Stephen Mitchell (Electron Microscope Facilities, King’s Buildings). We would like to acknowledge Natalie Ring for providing guidance on ONT sequencing and analysis.

---

**S1 Fig. Characterisation of phage LUC4.**

LUC4 is the most virulent phage in our collection that can target EC958. (A) Transmission electron microscopy reveals that LUC4 has the morphology of a myovirus. (B) The genome of phage LUC4 is 163,734 bp long with 49 genes involved in DNA, RNA and nucleotide metabolism, 11 tRNA genes, 10 transcriptional regulation genes, 29 genes encoding the tail, 3 genes encoding the connector proteins, 16 genes involved in the structure of the head and the packaging of the genome, 10 genes involved in the host takeover and auxiliary metabolism, and 7 genes involved in lysis. No virulence, AMR or lysogenic genes were found.

**S2 Fig. One-step growth curve of phage LUC4.**

A latent period of approximately 25 minutes and a burst size of 78 viral particles per infected cell were measured with a one-step growth curve of LUC4 with EC958 in AU.

**S3 Fig. OmpC mutations in fixed LUC4 mutants, complementation, fitness costs and infection dynamics of Chap1.**

**(A)** Fixed-resistance is driven by mutations and loss of function of the phage receptor OmpC: Genome sequence alignments from long-read sequence data illustrate diverse genomic rearrangements including large scale deletions, insertions and inversions, that result in the loss of OmpC or OmpR. **(B)** Expression of EC958 OmpC in ClearColi® confers susceptibility to phage LUC4. The *ompC* gene from EC958 was cloned into the pWKS30 vector and transformed into *E. coli* strain ClearColi®, which lacks native OmpC expression. Upon induction, ClearColi®+pEC958ompC became susceptible to phage LUC4 in both solid and liquid infection assays, confirming OmpC as the LUC4 receptor. **(C)** Competitive fitness of OmpC variants ECRA304.8 and ECRA544.4 relative to wild-type (WT) EC958. To test whether phage-resistant variants carried a fitness cost, competition assays were performed. Relative fitness of the two fixed-resistant variants compared to WT EC958 after 24 hours of co-culture in LB medium was <0.5. A fitness value >1.0 indicates a competitive advantage over WT EC958, while values <1.0 suggest reduced fitness. The fitness of the rifampicin-resistant variants was also measured in relation to the parent strain and showed a fitness cost that was less pronounced. **(D)** Phage Chap1 was isolated on a LUC4 resistant EC958 mutant. Graphs show growth dynamics of EC958 with LUC4, Chap1 and LUC4/Chap1 combined in LB, AU and PU. Data shown is one representative biological replicate, with growth curves displaying the mean ± SD of three technical replicates. Three independent biological replicates were performed.

**S4 Fig. Impact of capsule and colonic acid on phage infection and depolymerase active phage.**

**(A)** Phage scores (calculated with AUC data of with / without phage of a panel of 31 phage in AU. EC958 WT is wildtype strain, EC958 CAP is EC958ΔkpsD and EC958 CAP_CA is EC958 ΔkpsDΔwcaF. **(B)** Plaque morphology of four phage (LUC4, CHAP1, Phi3 and TapS) on host strain EC958 showing depolymerase halos for Phi3 and TapS.

**S5 Fig. Biolog analysis.** Phenotypic analysis of EC958 growth in presence (red) and absence (green) of phage LUC4 with different sole carbon sources using Biolog PM01 plate over a 48 hour incubation. Each carbon source is annotated in the appropriate panel.

**S1 Table. Variant calling with Snippy.**

Fixed-resistant variants were Illumina-sequenced. The paired reads were analysed with Snippy, which revealed 14 variants with SNPs or small INDELs disrupting *ompC*, *ompR*, *envZ* or the intergenic region immediately upstream from *ompC*, suggesting a loss of function of the phage receptor: *ompC*. No SNP or INDEL could be detected with Snippy in 16 of the samples. Two variants had SNPs in unknown or unrelated genes (*pstB* and *ruvB*).

**S2 Table. Sequence of the synthetically produced DNA flanking regions for *kpsD* and *luxS* knockouts.**

## Notes

### Competing Interest Statement

The authors have declared no competing interest.

